# Peripheral blood cell-type and sex-specific signatures of alcohol misuse revealed by single-cell transcriptomics

**DOI:** 10.1101/2025.05.21.655347

**Authors:** Amol C. Shetty, Angela Verma, John Sivinski, X Fan, Carolyn Doty, Brooke Hollander, Daniel Roche, Chamindi Seneviratne

**Author notes:** co-senior authors (Daniel Roche, Chamindi Seneviratne). **Corresponding Authors:** Daniel J.O. Roche, Ph.D., Assistant Professor, University of Maryland Baltimore, PO Box 21247, Baltimore, MD 21228, United States of America. Chamindi Seneviratne, Ph.D., Adjunct Assistant Professor, University of Maryland Baltimore, Baltimore, MD, 21201, USA.

## Abstract

Alcohol use disorder (AUD) is a complex condition with diverse molecular underpinnings. Chronic alcohol exposure is associated with alterations in both innate and adaptive immune cell populations that in turn contribute to AUD susceptibility and severity, suggesting a bidirectional gene-environment relationship. Yet, most clinical AUD studies of this system have relied on indirect measurement of immune function and/or assessment of only a small number of cell types or immune markers, approaches which cannot capture the immune system’s inherent complexity and cellular heterogeneity. Therefore, in order to better characterize immune dysregulation in AUD, the goal of this study was to use single cell RNA sequencing (scRNA-seq) in peripheral blood mononuclear cells (PBMCs) to compare immune cell proportions and differential gene expression between individuals with AUD and healthy controls. Our findings highlight a distinct disproportionality of lymphocytes and monocytes in individuals with AUD and healthy controls as well as sex and dose-dependent alterations immune cell functional characteristics. These peripheral blood cell-type proportions and dose-dependent signatures of alcohol exposure may aid in developing more targeted and effective pharmacological interventions for AUD. Results also highlight the importance in including sex as a biological variable in AUD research, particularly when examining immune function.

## INTRODUCTION

Alcohol use disorder (AUD) is a prevalent and costly disease for which current pharmacological treatments are only moderately effective.^1, 2^ Characterizing the biological mechanisms of AUD is a priority in order to identify novel pharmacological targets and develop more efficacious medications.^3^ Translational evidence supports the immune system as being one such target, as alterations in immune function may play a key role in disease progression.^4^ Yet, the immune system is dynamic, multi-cellular, and multi-organ, spanning peripheral and central nervous systems. Most clinical AUD studies of this system have relied on indirect measurement of immune function and/or assessment of only a small number of cell types or immune markers, approaches which cannot capture the immune system’s inherent complexity and cellular heterogeneity. Therefore, in order to better characterize immune dysregulation in AUD, the goal of this study was to use single cell RNA sequencing (scRNA-seq) in peripheral blood mononuclear cells (PBMCs) to compare immune cell proportions and differential gene expression between individuals with AUD and healthy controls.

Chronic, heavy alcohol exposure may produce systemic dysregulation in both the innate and adaptive immune systems.^4, 5^ Dysfunction in these systems appear to play a role in the acute effects of alcohol,^6, 7^ motivation to seek alcohol,^8–11^ and AUD-associated organ damage and susceptibility to infection.^12–15^ Despite earlier interpretations suggesting that AUD was associated with a sustained pro-inflammatory state driven by systemic innate immune system activation, a more complete, nuanced perspective has developed indicating the innate and adaptive immune systems each go through dynamic periods of activation and inactivation, with specific cell types and signaling pathways being up- or down-regulated as a function of drinking patterns and disease state. ^7, 16–18^ Most human AUD studies to date have studied immune function in the brain^7, 16, 19–21^ or by measurement of peripheral cytokine levels.^22, 23^ Thus, the cell-type-specific transcriptional changes that occur in the immune system of individuals with AUD have not been well characterized, potentially limiting our ability to identify immune-related pharmacological targets.

Single-cell RNA sequencing (scRNA-seq) has advanced understanding of the cellular composition of the human brain^24–26^ and molecular basis of psychiatric diseases^27, 28^ by enabling the simultaneous analysis of gene expression in thousands of individual cells. This technology allows identification of distinct cell populations and their specific transcriptional changes in complex tissues and fluids like the brain, liver, or blood, potentially revealing cell-specific molecular mechanisms masked in traditional analyses. Most studies in substance use disorder (SUD) and AUD have utilized Single-nucleus RNA sequencing in post-mortem brain samples or brain organoids.^27, 29, 30^ Relatively few studies have used scRNA-seq to identify changes in peripheral gene expression *in vivo* in individuals with AUD. Of those that have, all studies in AUD that employed scRNA-seq in peripheral blood mononuclear cells (PBMCs) to examine cell specific changes associated with the disorder have been in populations with liver disease of varied severity.^31–35^ As liver disease is associated with complex immune dysfunction independent of the influence of chronic alcohol exposure,^36, 37^ in order to understand the peripheral molecular mechanisms of alcohol-associated immune dysregulation, studies in individuals with AUD but without liver disease are needed. Thus, the present study used scRNA-seq in PBMCs in individuals with AUD without liver disease and healthy controls to investigate AUD and the dose-dependent effects of alcohol peripheral immune cell populations.

## MATERIALS AND METHODS

### Study design Participants

N=8 blood samples were collected at baseline from individuals with AUD (n=4) and healthy controls (n=4) who were participating in a randomized controlled medication trial (NCT04210713). The parent study and the methods described in this manuscript were approved by the University of Maryland Baltimore Institutional Review Board and conducted in accordance with the Declaration of Helsinki. Full description of recruitment, methods, and eligibility criteria are listed in the Supplemental Methods.

In brief, participants were recruited from the general community via social media advertisements. Interested participants completed a phone screen and, if initially eligible, were invited to an in-person screening session to further assess eligibility for the parent trial. The screening session began with a breath alcohol concentration test (0.00 g/dl required) and written informed consent. Participants then completed questionnaires and interviews about their medical history, mental health, sleep, and substance use. Eligible participants were randomized to placebo or minocycline and then attended a pre-medication baseline session that included blood, saliva, and stool sample collections, behavioral testing, and neuroimaging. Participants arrived at the baseline session at 9:30 AM, completed a breath alcohol concentration test (0.00 g/dl required), and then completed blood draws for the samples analyzed in this study.

### PBMC sample preparation

PBMCs were isolated from individuals with AUD and controls and immediately cryopreserved.

### RNA sequencing

#### scRNA-seq

A total of 9 million cells were loaded into each well of a chromium microfluidics controller (10X Genomics) to target 10,000 cells per sample. Sequencing libraries were generated using the Chromium Next GEM Single Cell 3’ Reagent Kits v3.1. Sequencing targeted 50,000 reads per cell. Samples were sequenced using Illumina paired-end 100bp sequencing (NovaSeq6000 S4 Lane - 2250M read pairs, 450 Gbp).

### Data analysis

To ensure high-quality cells were utilized for downstream single-cell analyses, we selected cells with a minimum of 100 genes and a maximum of 6,500 genes expressed and retained genes that were expressed in at least 3 cells. Since cells are often stressed, damaged, and undergo apoptosis during sample preparation, we retained cells with mitochondrial gene proportions below 25%. Using these thresholds, we retained ∼38,000 high-quality cells ranging from ∼2,850 to ∼6,650 cells per sample. ∼38,000 high-quality cells from 8 PBMC samples were analyzed using Seurat (v4.3.0)^38^for normalization of the gene expression data, identification of variable gene features and dimensionality reduction using Principal Component Analysis (PCA) and Uniform Manifold Approximation and Projection (UMAP). Cells across multiple samples were integrated using integration anchors identified using the canonical correlation analysis method to remove potential batch effects, followed by data scaling, and dimensionality reduction of the integrated dataset.

Post-integration the cells were grouped into 5 major immune cell clusters that were further sub-clustered into 25 different immune cell clusters using Seurat (v4.3.0). Cell-cluster specific marker genes were identified for each cluster that were then compared to marker genes curated from CellMarker databases^39, 40^ and relevant cardiovascular-related literature. Using these cell annotations, we labeled the 25 cell clusters to identify thirteen T lymphoid immune cell sub-clusters, four B lymphoid immune cell sub-clusters, and seven myeloid immune cell clusters wherein each cluster consisted of ∼30 – ∼5,500 cells per cluster. Differential expressed genes (DEGs) were identified for each of the annotated clusters separately for comparisons between cells belonging to two different conditions, namely AUD vs sex-matched controls, high or low DpDD vs sex-matched controls, and different TD vs sex-matched controls. Default methods available through Seurat (v4.3.0) utilizing the Wilcoxon Rank Sum test were used to identify DEGs with a minimum log2-scaled fold-change of 0.25 and for genes expressed in at least 25% of cells in either of the two groups. Over representation analysis method was then used to assess and evaluate the overlap of the DEGs with known functional gene sets curated from Geno Ontology databases,^41^ Hallmark,^42^ KEGG,^43^ and REACTOME^44^ Pathways which enables the biological interpretation and functional significance of the DEGs identified for different cell clusters between different conditions.

## RESULTS

### Sample Characteristics

Table 1 presents sample demographics, alcohol and other substance use, AUD and other psychiatric diagnoses, and medication use and medical conditions.

**Table 1.**
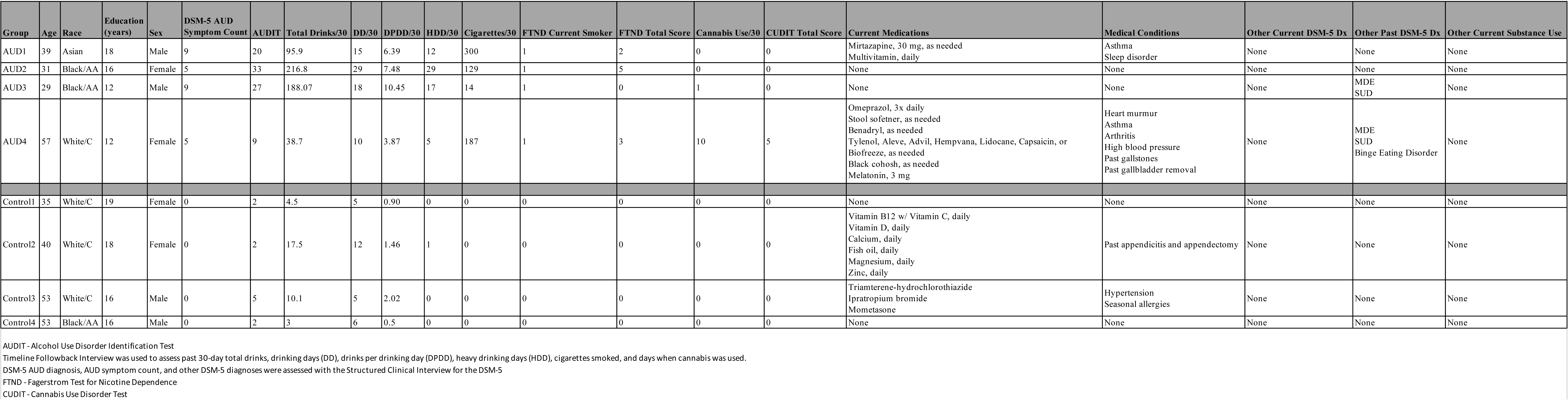

### Major lymphoid and myeloid cell types identified in individuals with AUD and controls

To obtain cell type-specific transcriptional profiles in AUD, we performed single-cell RNA-sequencing (scRNA-seq) analysis on the peripheral blood collected from individuals with AUD and controls. In total ∼38,000 high-quality cells were assessed across 4 individuals with AUD and 4 controls (ranging from ∼2,850 to ∼6,650 cells) and grouped into 25 clusters which expressed classic immune cell type markers (Figure 1A).

**Fig. 1.**
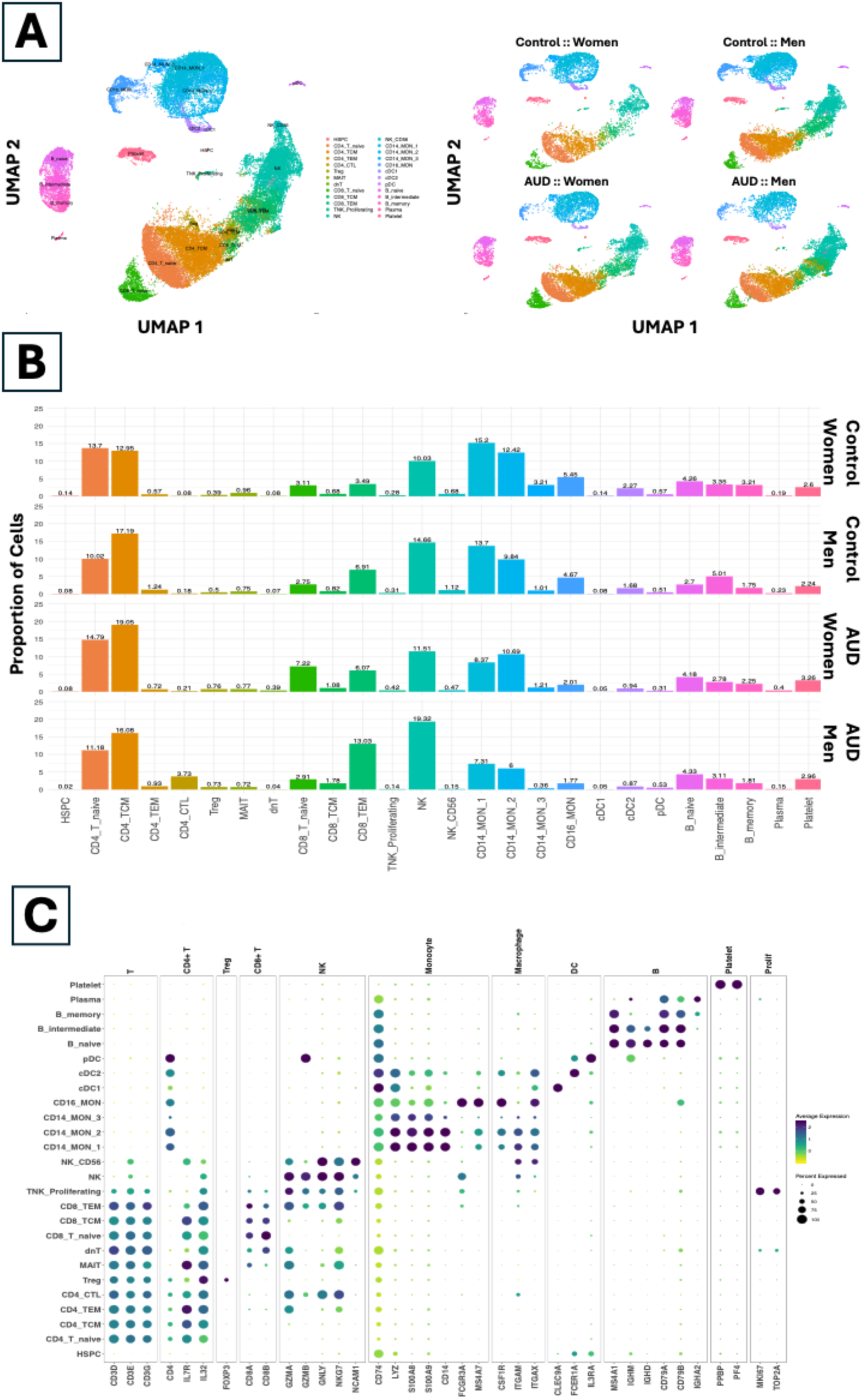
scRNASeq cell clustering and annotation. The scRNASeq dataset consisted of **(A)** ∼38,000 high-quality cells from PBMC samples collected from women and men with AUD and controls that were grouped into 25 different immune cell clusters. **(B)** The cell proportions for each cluster varied between individuals in AUD and controls and **(C)** the cell clusters were annotated based on the expression of known immune cell markers illustrated as a dotplot where the color gradient represents the scaled expression, and the size of the dot represents the percent of cells expressing the specific immune cell marker.

After cell clustering and annotation, we identified four CD4^+^ T lymphocyte (CD4_T: 28% Control, 33% AUD) clusters, three CD8^+^ T lymphocyte (CD8_T: 9% Control, 16% AUD) clusters, two natural killer (NK: 14% Control, 16% AUD) clusters, three B lymphocyte (10% Control, 9% AUD) clusters, four monocyte (MON: 32% Control, 19% AUD) clusters, three dendritic cell (DC: 3% Control, 1% AUD) clusters and other cell types such as regulatory T cells (Treg: <1% in both Control and AUD), mucosal-associated invariant T cells (MAIT: <1% in both Control and AUD), plasma cells (<1% in both Control and AUD), proliferating T and NK cells (<1% in both Control and AUD), stem cells (<1% in both Control and AUD), and platelets (2% Control, 3% AUD).

In individuals with AUD, we observed higher proportions of CD8^+^ T memory cells, NK cells, and CD4^+^ T cytotoxic cells and lower proportions of CD14^+^ and CD16^+^ monocyte cells (Figure 1B). When assessed separately for men and women with AUD, we observed higher proportions of CD4^+^ T memory, naïve CD8^+^ T, and CD8^+^ T memory cells in women with AUD, higher proportions of CD8^+^ T memory cells, NK cells, and CD4^+^ T cytotoxic cells in men with AUD, and lower proportions of CD14^+^ and CD16^+^ monocyte cells in both men and women with AUD (Figure 1B).

Gene expression profiles of canonical markers of lymphoid and myeloid cells confirmed the cell annotations (Figure 1C), and cell type-specific markers identified molecular differences between sub-clusters of a particular cell type. CD4^+^ T effector memory cells showed overexpression of granzyme K (*GZMK*) while CD4^+^ T cytotoxic cells showed overexpression of granzyme H (*GZMH*). Among the CD8^+^ T cell clusters, we observed an enrichment of naïve CD73^+^ CD8^+^ T cells in women with AUD which could exert and immunosuppressive effect and an enrichment of TRGC2^+^ CD8^+^ T effector memory cells, a distinct subset of γδ T cells, in men with AUD. Both men and women with AUD also show depletion of S100A^+^ CD14^+^ CD16^-^ monocytes and CDKN1C^+^ CD14^-^ CD16^+^ monocytes. The above results focused on cell type annotation and distribution support the diversity of cell types that are enriched/depleted in the peripheral blood from individuals with AUD compared to controls and the heterogeneity of lymphoid cell types enriched in men and women with AUD.

### Altered cell type-specific gene expression profiles between individuals with AUD and controls

Next, we compared the cell type specific transcriptional profiles between individuals with AUD and controls separately for men and women with AUD to identify genes dysregulated in individuals with AUD (Figure 2). In women with AUD, we identified IFI44L, IFIT1, IFIT3, and CENPK to be downregulated in CD4^+^ T memory cells and naïve CD4^+^ T cells that play an important role in the regulation of the immune response, inflammation, and cell division respectively (Figure 2A). We also identified down-regulation of IFI44L, IFI44, and MX1 and up-regulation of granzymes and fibroblast growth factors in CD8^+^ T effector memory cells that are known to play a role in immune response and cell growth. CD14^+^ monocytes in women with AUD showed up-regulation of RETN, NFXL1, and DLGAP2. Similar assessments in men with AUD showed MTRNR2L8, ARL17B, and CD27 to be down-regulated and up-regulation of CD226, ATP10A, and ITGB1 in CD8^+^ T effector memory cells (Figure 2B). These genes may play a role in regulating the immune system, transmembrane transportation, and cell-cell communication. We also identified up-regulation of multiple HLA genes, and SLC35F1 in NK cells that are known to play a role in regulating the immune response, activity of NK cells, and transmission of neuronal signals. In CD4^+^ cytotoxic T cells that were enriched in men with AUD only, we observed up-regulation of genes such as SLC15A4 and SLC35F1 that encode for solute carrier proteins, and LAIR2, a member of the immunoglobulin superfamily that are responsible for the transport of molecules across the cell membrane. CD14+ monocytes in men with AUD showed up-regulation of HLA-DRB5, SLC35F1, and CSF1R. More research is needed to confirm the role of these immune-related genes in AUD. Additionally in women and men with AUD, we observed a reduction of CD14^+^ monocytes that showed over-expression of interferon-stimulated genes (ISGs) such as IFIT1, IFIT2, and IFIT3, and intracellular protein transport and vesicle-mediated transport gene ARL17B that play a crucial role in the innate immune response. We also observed that cell type-specific dysregulation of some genes presented differently in women with AUD compared to men with AUD (Figure 2C).

**Fig. 2.**
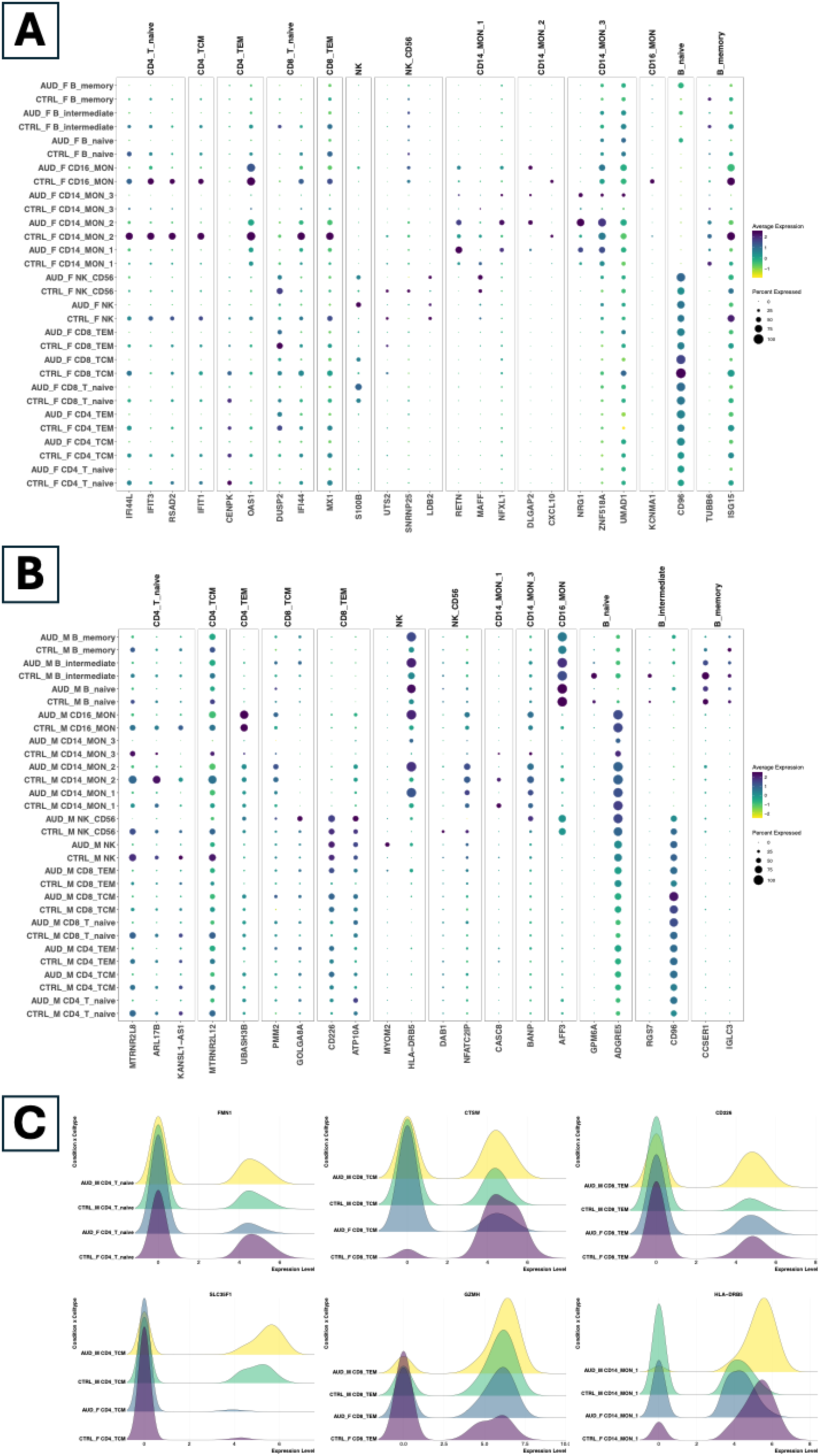
Differential gene expression between individuals with AUD compared to controls. Comparison of gene expression between individuals with AUD and controls identified several genes dysregulated within specific immune cell types in **(A)** women with AUD and **(B)** men with AUD. The gene expression of genes is illustrated as a dotplot where the color gradient represents the scaled expression, and the size of the dot represents the percent of cells expressing the specific gene. Cell type-specific gene expression also showed **(C)** differences in gene dysregulation between women with AUD and men with AUD that are illustrated as density plots of gene expression for a candidate gene within a specific cell type for individuals with AUD and controls stratified by sex.

### Altered cell type-specific gene expression profiles driven by higher drinks per drinking day (DpDD) in individuals with AUD

Given the cell type specific dysregulation identified in individuals with AUD and the sex-specific transcriptional differences, we further assessed the cell type specific transcriptional profiles between higher and lower drinks per drinking day (DpDD) in individuals with AUD, separately for men and women with AUD, to identify dysregulation associated with DpDD. The DpDD in individuals with AUD varied from 3.9 drinks to 10.5 drinks compared to a maximum of 2.0 drinks in controls. In women with AUD, we observed increased proportions of naïve CD4^+^ T, CD4^+^ T central memory, and naïve CD8^+^ T cells as DpDD increased and decreased proportions of CD14^+^ monocytes as DpDD increased (Figure 3A). In naïve CD4^+^ T, CD4^+^ T central memory, and naïve CD8^+^ T cells, we identified long non-coding RNA (lncRNA) SNHG31, a potassium channel protein KCNQ5, a T cell activation marker CD63, and a cytokine signaling gene PASK as immune markers associated with increasing DpDD in women with AUD. NK cells also showed over-expression of S100B, KLRC genes, and IL32 that are involved in NK stimulation and enhanced cytotoxic effect with increasing DpDD in women with AUD (Figure 3B). In CD14+ monocytes, RETN expression increased while HLA gene expression decreased with increasing DpDD in women with AUD. Similarly, in men with AUD, we observed increased proportions of CD4^+^ cytotoxic T, CD8^+^ T effector memory, and NK cells as DpDD increased and decreased proportions of CD14^+^ monocytes as DpDD increased (Figure 3A). CD4^+^ T central memory cells showed FKBP5, SMDT1, and NIBAN1 that play a role in immune regulation, mitochondrial function, and stress response/apoptosis to be associated with increasing DpDD in men with AUD. In CD4^+^ cytotoxic T cells, we found SLC35F1 over expressed in men with AUD with high DpDD. In CD8^+^ T effector memory cells, we identified FNDC3B that involved in cell adhesion, FKBP5 involved in immune response induced by corticosteroids, and LGALS1 involved in apoptosis as immune markers associated with increasing DpDD in men with AUD. Additionally, NK cells showed over-expression of CD52, GZMH, LGALS1, CRIP1, and S100A6 that are involved in NK cytotoxicity and cytokine production with increasing DpDD in men with AUD (Figure 3C).

**Fig. 3.**
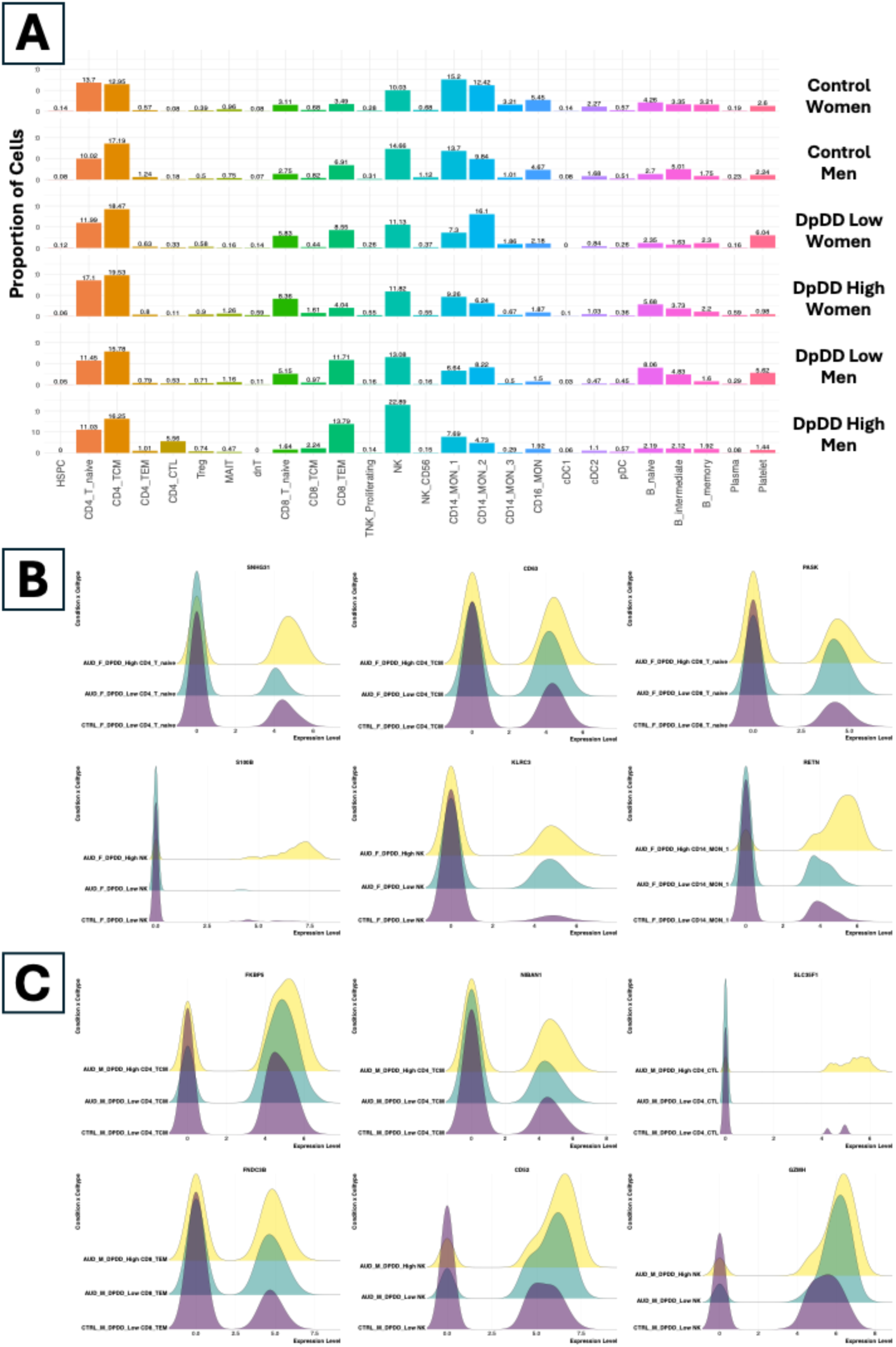
Differential gene expression between individuals with AUD stratified by DpDD in women and men. The cell proportions for each cluster **(A)** varied between women and men with AUD stratified by DpDD and controls. Cell type-specific gene expression also showed association of gene dysregulation with increasing DpDD in **(B)** women with AUD and **(C)** men with AUD that are illustrated as density plots of gene expression for a candidate gene within a specific cell type for women and men with AUD with varying DpDD and controls.

### Altered cell type-specific gene expression profiles driven by total drink (TD) consumed by individuals with AUD

Next, we assessed the cell type specific transcriptional profiles associated with varying levels of total drinks (TD) in individuals with AUD. The total drinks in individuals with AUD varied from 39 drinks to 217 drinks compared to a maximum of 17.5 drinks in controls. In individuals with AUD, as TD increases, we observe high proportions of naïve CD4^+^ T, CD4^+^ T central memory, naïve CD8^+^ T, CD8^+^ T effector memory, and NK cells (Figure 4A). In CD4^+^ T central memory cells, we observed higher expression of TMEM14C, CD63, and TRBC1 when compared to sex-matched controls as TD levels increased. In CD8^+^ T effector memory cells, we observed higher expression of ITM2A, MRPL51, TRMT112, and SF3B6 with increasing TD levels in individuals with AUD. In NK cells, we observed higher expression of DOCK5, DTNBP1, and TSPO as TD levels increased in individuals with AUD. (Figure 4B) All the above results confirm that the genetic basis of AUD is complex and involves multiple cell-types and genes, in addition to environmental and lifestyle factors.

**Fig. 4.**
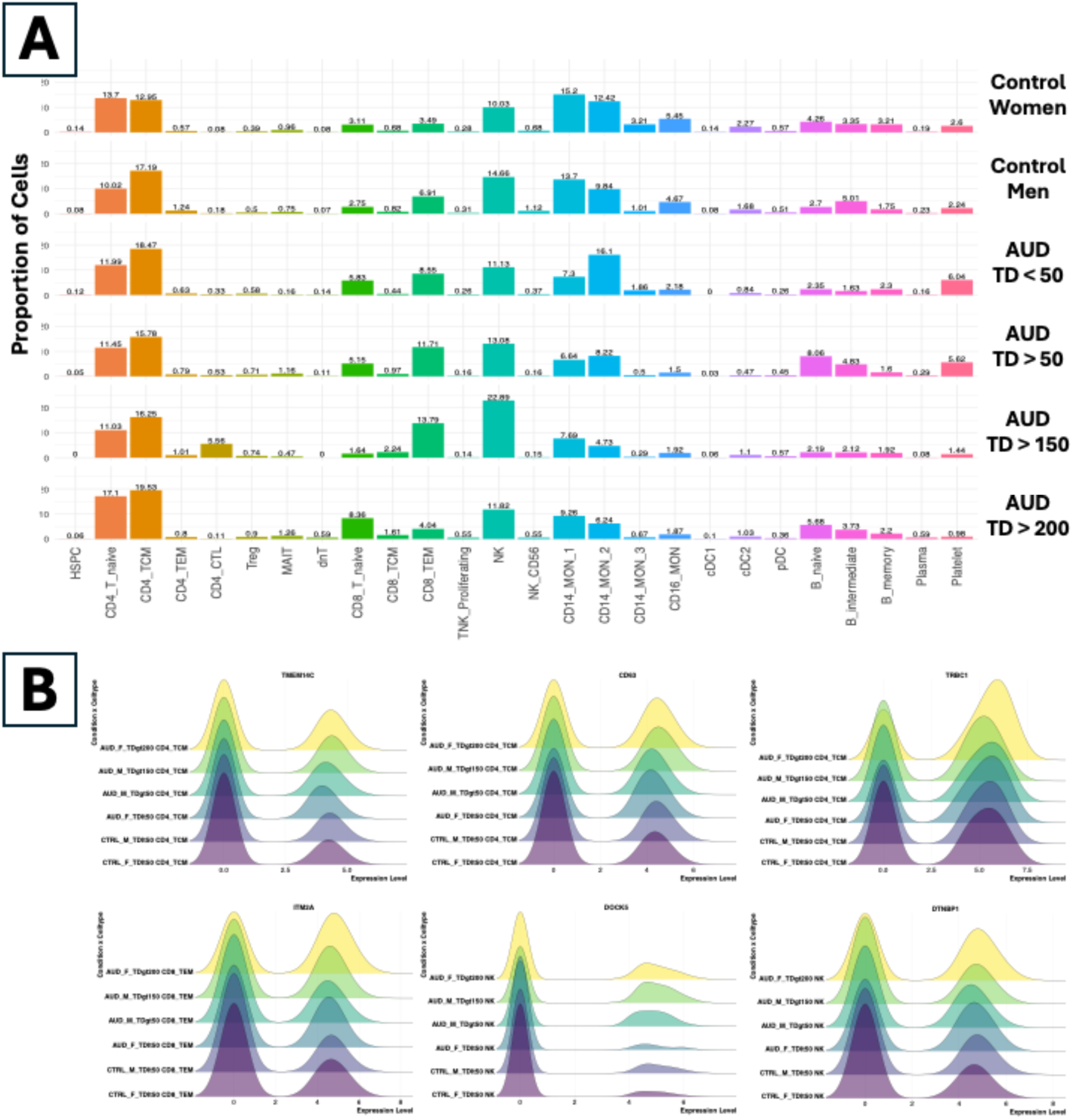
Differential gene expression between individuals with AUD stratified by TD . The cell proportions for each cluster **(A)** varied between women and men with AUD stratified by DpDD and controls. **(B)** Cell type-specific gene expression also showed association of gene dysregulation with increasing **TD in individuals** with AUD that are illustrated as density plots of gene expression for a candidate gene within a specific cell type for women and men with AUD with varying DpDD and controls.

## DISCUSSION

The primary goal of our study was to elucidate inherent complexity and cellular heterogeneity of immune cells in the blood of individuals with AUD using a single-cell transcriptomics approach. We aimed to identify how chronic alcohol exposure alters the function and communication of various immune cell types within the peripheral blood. Understanding these immune dysregulations could reveal novel biomarkers for AUD diagnosis as well as potential targets for pharmacologic intervention. Ultimately, this knowledge may be crucial for developing targeted immunomodulatory therapies to alleviate symptoms of chronic alcohol exposure and mitigate alcohol-related organ damage. In order to assess changes transcriptomic profiles associated with chronic alcohol use, we sequenced samples from individuals with AUD (n=4) and healthy controls (n=4) at the single-cell level. We found that AUD was associated with robust, dose dependent changes in the distribution of immune cell types as well as differential gene expression within these cells and that many of these results were moderated by sex.

Overall, we observed a heterogeneous population of both lymphoid and myeloid immune cells in both individuals with AUD and healthy controls. However, individuals with AUD, relative to controls, showed higher proportions of adaptive immune T lymphocytes (i.e., CD8^+^ T memory cells and CD4^+^ T cytotoxic cells) and innate immune NK cells as well as lower proportions of innate immune monocytes (i.e., CD14^+^ and CD16^+^ cells). Such findings align with prior work demonstrating that chronic alcohol exposure *in vitro* and *in vivo* affects the number and function of T-cells,^45^ in particular an acceleration in the proliferation, differentiation, and conversion of naïve T cells into memory T cells.^46–49^ Prior studies have shown increased NK cells in individuals with AALD,^50^ but ours may be the first to suggest an increase before diagnosed AALD. Further, acute and chronic alcohol exposure produces changes CD14+ and CD16+ cell number and function, specifically depletion in CD14^+^ CD16^-^ monocytes.^51^ Such shifts in the balance of innate and adaptive immune cell types has been theorized to underlie development of a decreased ability to recognize and fight infection in AUD.^45^

Transcriptional differences between individuals with AUD and healthy controls revealed several dysregulated genes in AUD that play an important role in regulation of the immune response, inflammation, cell division, and cell growth in multiple T lymphocyte populations. In individuals with AUD, we observed dose-dependent upregulation of several genes previously shown to involved in liver and gut cancer,^52–54^ AALD,^55^ and overall liver health.^56, 57^ In those with AUD, we observed down-regulation of genes that play a crucial role in the innate immune response, including interferon-stimulated genes (ISGs) and intracellular protein transport genes in CD14+ monocytes. Downregulated ISG and/or dysfunctional response to interferon has been previously documented individuals with AALD and heavy drinkers and may be an adaptation to chronic overactivation of interferon/TLR-mediation immune pathways.^58, 59^ Further assessing the impact of DpDD on cellular composition, we identified a blunted effect of alcohol dose on naive T lymphocytes in women with AUD and greater alcohol dose effects on cytotoxic T lymphocytes and natural killer immune cells in men with AUD. It is notable that expression of FK506-binding protein 51 (FKBP5) was positively associated with alcohol dose only men, as activity of this gene has been translationally linked to AUD and stress-related disorders,^60–62^ and inhibition of the FKBP5 protein is a promising intervention target for AUD.^63^ Lastly, we investigated the impact of increasing total drinks in individuals with AUD and showed higher expression of genes involved in T-cell activation and cytotoxic activity. Further studies are warranted to characterize the sex and dose dependent functions of these genes and signaling pathways in AUD.

The study has several strengths, including using a deeply phenotyped cohort with no history of AALD or immune-related disorders. Study limitations include a smaller sample size due to specimen availability and the presence of other clinical comorbidities, including nicotine dependence which has shown to affect immune cells.^64^ However, our findings serve as hypothesis generating insights and recognize the need to better understand the cellular heterogeneity and activity composed of different lymphoid and myeloid immune cells in individuals with AUD. Our results will guide future cell type-specific studies to derive biomarkers for chronic alcohol exposure in individuals with AUD. In conclusion, our exploratory analysis at a cellular level has identified several peripheral blood cell-type and sex-specific signatures of alcohol misuse in individuals with AUD that are driven by distinct dysregulation of the innate and adaptive immune system.

## List of Supplementary Materials

### Supplementary Methods

Table S1: Differential expression for genes for individual celltypes associated with AUD.

Table S2: Reactome pathways enriched by differential expressed genes associated with AUD.

Table S3: Differential expression for genes for individual celltypes associated with DpDD.

Table S4: Reactome pathways enriched by differential expressed genes associated with DpDD.

Table S5: Differential expression for genes for individual celltypes associated with TD.

Table S6: Reactome pathways enriched by differential expressed genes associated with TD.

### Funding

National Institute on Alcohol Abuse and Alcoholism grant to DJOR K01AA026005

## Notes

### Competing Interest Statement

The authors have declared no competing interest.

